# Computational study of the effect of hypoxia on cancer response to radiation treatment

**DOI:** 10.1101/2020.10.21.348474

**Authors:** T. Korhonen, J. H. Lagerlöf, A. Muntean

**Affiliations:** Department of Mathematics and Computer Science, Karlstad University, Sweden; Department of Medical Physics, Karlstad Central Hospital, Karlstad, Sweden; Department of Medical Physics, Faculty of Medicine and Health, Örebro University, Örebro, Sweden

## Abstract

We perform a computational study of the propagation of the oxygen concentration within a two-dimensional slice of a heterogeneous tumour region where the position and shape of the blood vessels are known. Exploiting the parameters space, we explore which effect is noticeable what concerns the formation of hypoxic zones. We use this information to anticipate a patient-specific radiation treatment with controlled response of the cancer growth.

## Introduction

It has long been known that the lack of oxygen in cells (hypoxia) greatly can hinder the effectiveness of a radiation treatment of cancer patients [2]. The concept of hypoxic/normoxic fractions is insufficient for predicting radiation treatment outcomes [3]. A good control of the tissue oxygenation is desired if one targets an excellent radiation of localised cancer regions with a minimal damage on the neighbouring tissue. This is the rationale for attempting to model, in detail, the tumour oxygenation. It is worth noting that the literature on the modeling and simulation of cancer radio- and chemo-therapies is large and keeps growing rather fast. At this stage, we only mention [4, 6, 8] and references cited therein.

The amount of oxygen present locally in a tumour can be measured directly by invasive methods (e.g. via sticking a needle). Pointwise indications can be obtained also via imaging techniques, see e.g. [10] and references cited therein. In this context, we support a non-invasive investigation of hypoxia by combining knowledge of the existing vasculature and numerical estimations of the oxygen transport and metabolism.

From the modeling perspective, the key is understanding the transport of oxygen through the blood vessels and through the surrounding heterogeneous environment. Within this framework we focus the attention on the transport of oxygen outgoing from the blood vessels travelling through both healthy and cancerous regions. The dynamics appears to be simple: oxygen is supplied by blood vessels from which it is then diffused in an environment where cells consume it. That is, as a first attempt, we can formulate a macroscopic diffusion-reaction model for the diffusion and consumption of oxygen in any given region when the arrangement of blood vessels and reasonable assumptions can be stated on the structure of the effective diffusion coefficient of oxygen as well as on the effective production rates by consumption.

Trusting previous works in this field (compare e.g. [9] and references cited therein), our model equations are as listed in (1). We denote by Ω the spatial region which includes the network of blood vessels and where the oxygen diffusion takes place during a time interval (0, *T*_*fin*_). Here *T*_*fin*_ is the final observation time of the overall process. Our modeling and computations are supposed to be valid until an *a priori* fixed time instance *T* ≤ *T*_*fin*_ (both *T* and *T*_*fin*_ are supposed to be close to the moment when the stationary state is reached). We denote by *K* = *K*(*x*, *t*) the mass concentration of oxygen distributed in the region Ω \ *V*, where by *V* we denote the region occupied by the blood vessels. Here *x* ∈ Ω \ *V* is the space variable while *t* ∈ (0, *T*) is the time variable.

Our problem boils down to computing the oxygen concentration *K*(*x*, *t*) satisfying the following model equations:

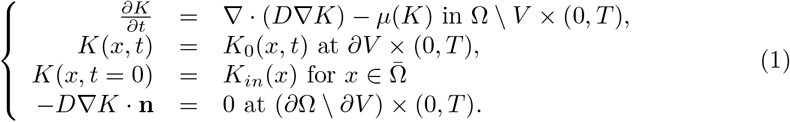

In this context, **n** is the outer normal vector to *∂V*, *D* = 2000*μm*^2^/*s* is an averaged diffusion coefficient for oxygen in water, *K*_0_ = 40mmHg^1^ is the supply of oxygen by the blood vessels^2^. Along the lines of [7], e.g., we assume for the production term in (1) the Michaelis-Menten consumption structure:

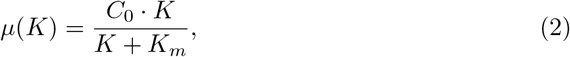

where *C*_0_ = 15mmHg/s and *K*_*m*_ = 1mmHg.

Interestingly, the model (1) is essentially a 3D-one; no correct reduction to a 2D setting can be made simply because the network of blood vessels is too complex. It is inherently a 3D object with no special structure (unless for particular cases). Hence, a dimension reduction exercise from 3D to 2D cannot be done in general without introducing too much uncontrolled errors in interpreting the geometry of the vasculature. As far as we see, treating such a problem correctly is a challenging task that arises in more general contexts when transport and storage in complex networks interact with the bulk environment. Instead of proposing *ad hoc* dimension reduction attempts, we study in this framework a cross-section in the tumour region. Such 2D domain is heterogeneous in the sense that it is composed of two sub-domains – a connected region where diffusion and production of oxygen takes place as well as a number of disconnected tiny regions where the blood vessels are located. These tiny regions are assumed to have a disk shape with eventually they can have a different radius from place to place as well as with different level of activity^3^. The reason for doing so is twofold:

i. We build an integrated tool for the fast detection *in silico* of hypoxic regions in tumours. This is a preparation step for handling a truly 3D setting.
ii. Currently, we notice the lack of an efficient mesh generation process from slice figures to mesh that could be imported into the computational platform FEniCS; see [1] for the users manual.

The main question we ask is: (Q) *Given the network as well as level of efficiency of blood vessels in a tumour, can one detect the oxygen diffusion regions where hypoxia is likely to be present?*

We address this question via a simulation study of the model 1 when the spatial distribution of blood vessels (occupying the region *V*) is provided. Our results indicate that the main challenge is to capture in detail the correct heterogeneous time and space distribution of the oxygen concentration. Once available in real time, this information can be used further to fine-tune a patient-specific radiation treatment.

## Materials and methods

### Example of a network of blood vessels

This section contains information on the topology of our reference network of blood vessels. The vessel images used in the simulations are microscopic photographs of 12 *μ*m slices of pancreatic islet cell carcinoma in transgenic RIP-Tag mice. This is a well vascularised tumour form. The endothelial cells (vessel walls) were dyed green and cell nuclei blue. The data is from Lagerlöf et al [7].

By visual inspection of our available images on blood vessels, it seems that most of the vessels are anyhow running perpendicular to the image surface and hence the effect of vessels running ‘along image surface’ is supposed to be small. This allows us to process good quality cross sections.

### Generation of the computational mesh

We employ GMSH [5] to generate meshes that accurately capture the underlying vessel structure from the 2D-slice images. We first obtain the vessels by thresholding the green channel of the slice image so that all pixels with green channel having value greater than 32 are considered to belong to a vessel. Subsequently employing image processing (now in the binary images vessel/not-vessel) we detect the vessel edges and further obtain contours along those edges which results *∂*Ω in Eq. (1), see Fig. 1. The vessels touching the image edges are closed and the domain is extended slightly so that vessel boundaries do not touch the boundary of the simulation domain. We can then supply *∂*Ω to GMSH and request a mesh tailored for the domain Ω \ *V*; see Fig. 1 (Right).

**Fig 1.**
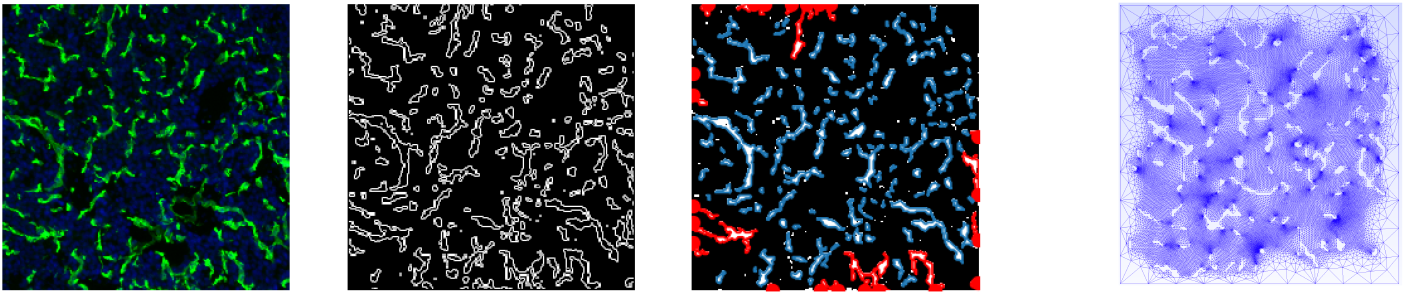
From image to simulation domain mesh. Left: Image processing is used to pick up the edge contours of the vessels (dyed green) from the original images. Right: GMSH generated mesh after being supplied with the vessel contours and domain boundary. The vessel edges are labeled so that each vessel can be uniquely picked up and assigned with a vessel surface oxygenation *K*_0_ using the Dirichlet boundary conditions.

### FEM approximation of the oxygen spatial distribution

The model equations admit a representation of the oxygen concentration in terms of an abstract Green function. Since the diffusion region Ω \ *V* has a complicated geometric structure, the structure of the Green function is not accessible. Instead, we use a finite element discretisation to approximate the model equations. These numerical approximations will be used together with the above image processing of the tumours’ vasculature to detect the hypoxic regions and give advice on scheduling irradiation treatments.

Firstly, we homogenise the Dirichlet boundary condition acting on the boundary of the region *V* occupied by the blood vessels, i.e. we set

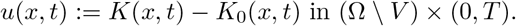

We rephrase the model equations as:

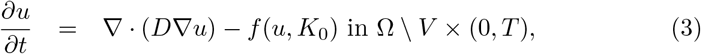

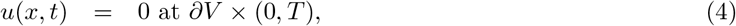

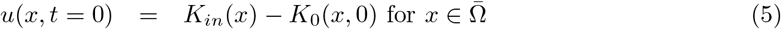

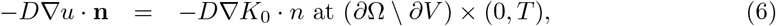

where the production term in (3) is given by

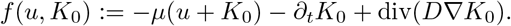

Based on (5), we observe that it makes sense to consider a parameter regime so that *K*_*in*_ − *K*_0_ ≥ 0. In this context, we are going to choose a constant value for *K*_0_ to represent the contact with a “constant reservoir of oxygen” coming from the blood vessels. However, the constant *K*_0_ can vary between vessels as later indicated.

The corresponding weak formulation reads: Find *u* such that for all suitable test functions *φ* the following integral identity holds:

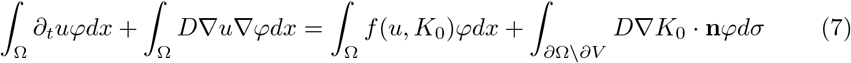

## Results

We use the tailor-made mesh produced with the help of GMSH to discretise efficiently our model equations. To handle (1) with corresponding initial and boundary conditions, we use the open source computational platform FEniCS; see for details [1]. The reference set of parameters is taken from [7, 9]. Linearising and transforming to weak form yields the expressions

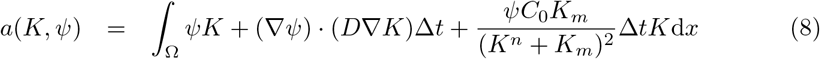

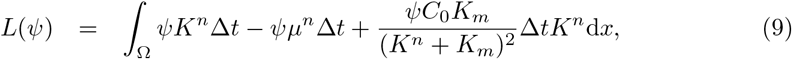

where *ψ* is test function, *K*^*n*^ is the oxygen concentration evaluated at the previous time step, while *μ*^*n*^ = *μ*(*K*^*n*^). To compute the correct approximate evolution of the overall process, we need to ensure that (*K* − *K*^*n*^) =: *δK* << *K* + *K*_*m*_ as this was used to linearise *μ*(*K*) from Eq. 2. We use Δ*t* = 20*μs*, which guarantees that the condition for *δK* is satisfied. GMSH takes care in the automatic generation of the adaptive mesh, which accounts for the precise distribution of the blood vessels.

The main output of our simulations consists of oxygen distribution profiles within the tissue. As post-processing work, we evaluate the local oxygen density (we refer to it in terms of histograms) as well as the expected irradiation response.

### Oxygen distribution in tissue

Due to e.g. vasomotion, the set of active vessels, i.e., those delivering oxygen to tissue as well as their oxygenation *K*_0_ can vary. For a given slice in the tumour we model this variation by randomly sampling a percentage (*active vessel percentage* takes values from [15, 30, 50, 75, 100]) of vessels to be ‘active’, that is, they have a surface *K*_0_ set to non-zero value. Moreover, for any given active vessel there is a probability (1/3) that the *K*_0_ is set to high value (50 mmHg) to create variation in the vessel surface oxygen concentrations.

We let the oxygen diffuse through the tissue while keeping the concentration at the vessel surface constant. After the steady state is reached, i.e. (in practical terms) the maximum oxygen concentration change between two time steps in every cell is less than 0.01%, we extract oxygenation distributions Figs. 2-4. This process is repeated over 10 times with random sampling of active vessels in each run. To obtain the histograms, we average over the 10 runs and further also extract standard deviation for robustness estimate. The chosen percentage of active vessels has a major impact on the number and area of hypoxic regions formed in the tissue; see the dark blue zones in Fig. 2.

**Fig 2.**
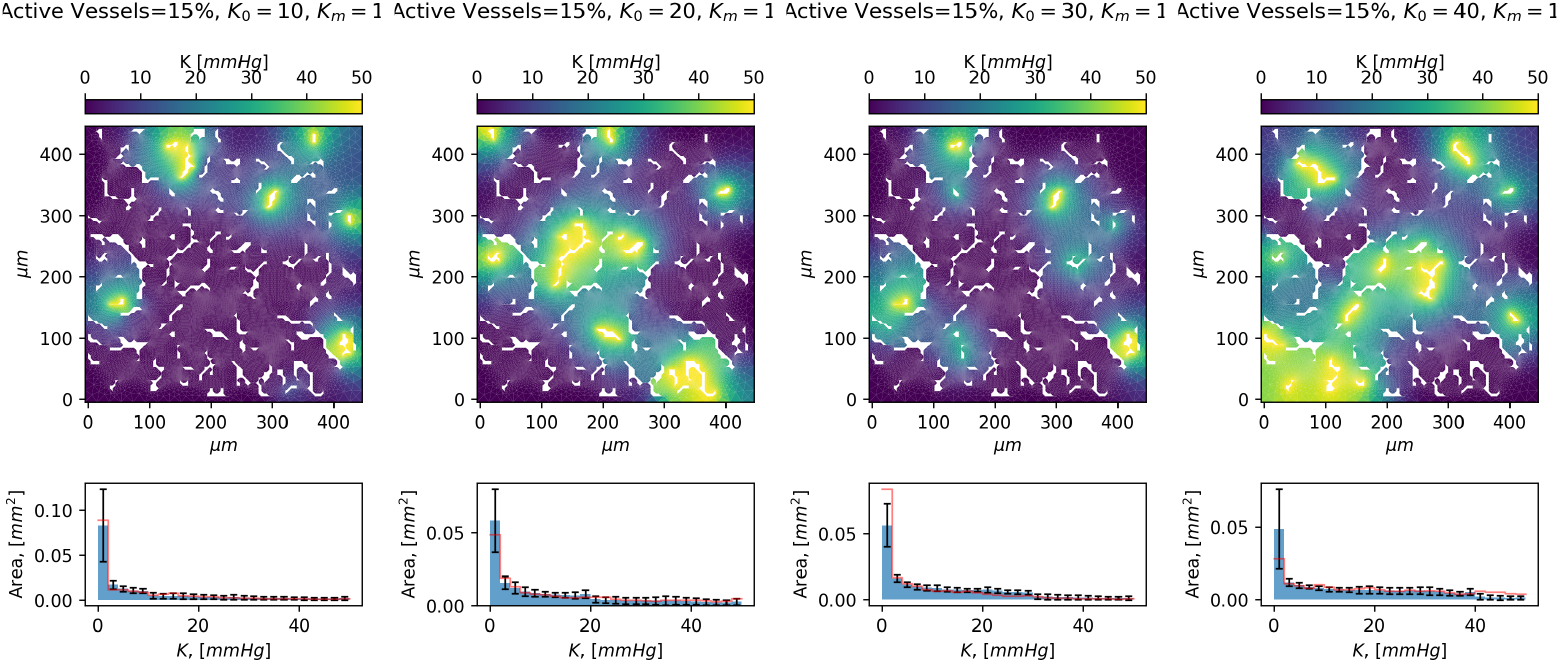
Top: Example oxygen concentration heat maps. Active vessels 15% and the surface *K*_0_ are from left to right: [10, 20, 30, 40]mmHg. Bottom: The mean oxygen concentration histogram is displayed along with standard deviation over the 10 runs.

**Fig 3.**
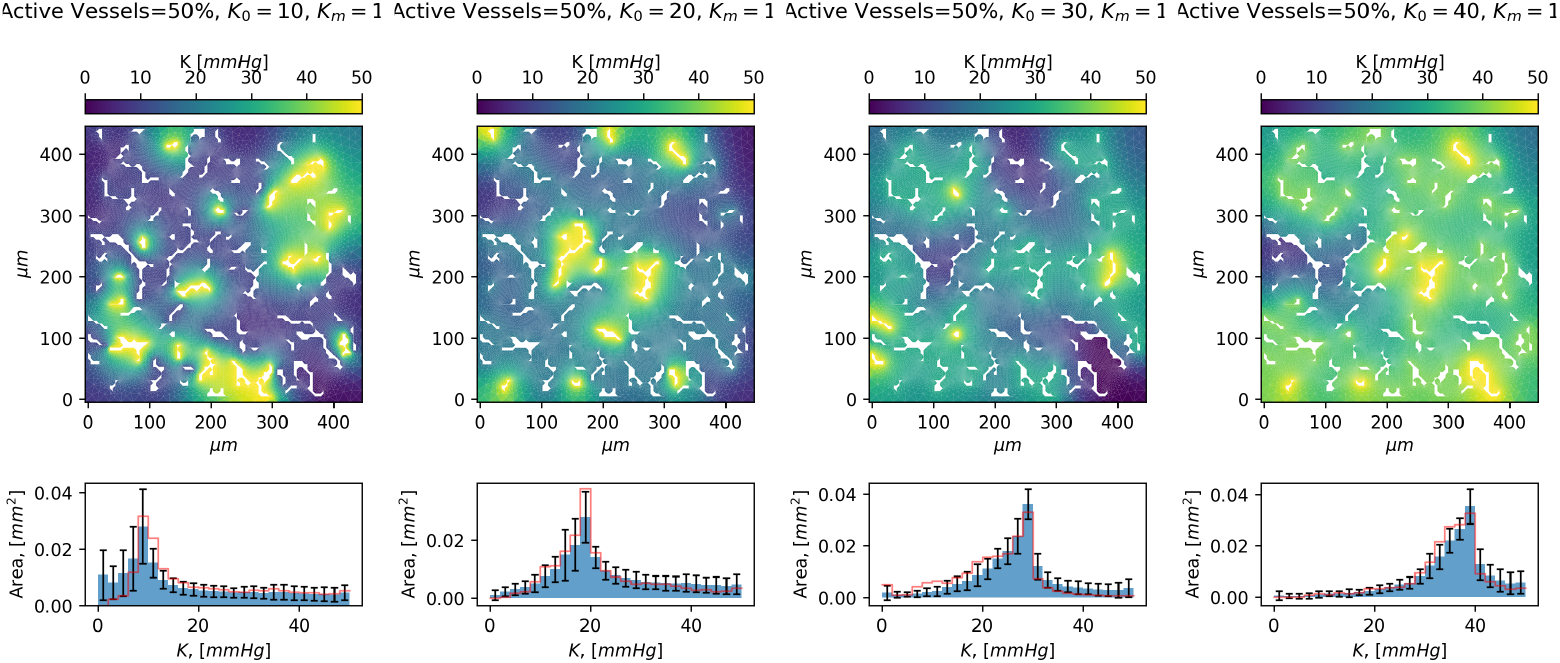
Top: Example oxygen concentration heat maps. Active vessels 50% and the surface *K*_0_ are from left to right: [10, 20, 30, 40]mmHg. Bottom: The mean oxygen concentration histogram is displayed along with standard deviation over the 10 runs.

**Fig 4.**
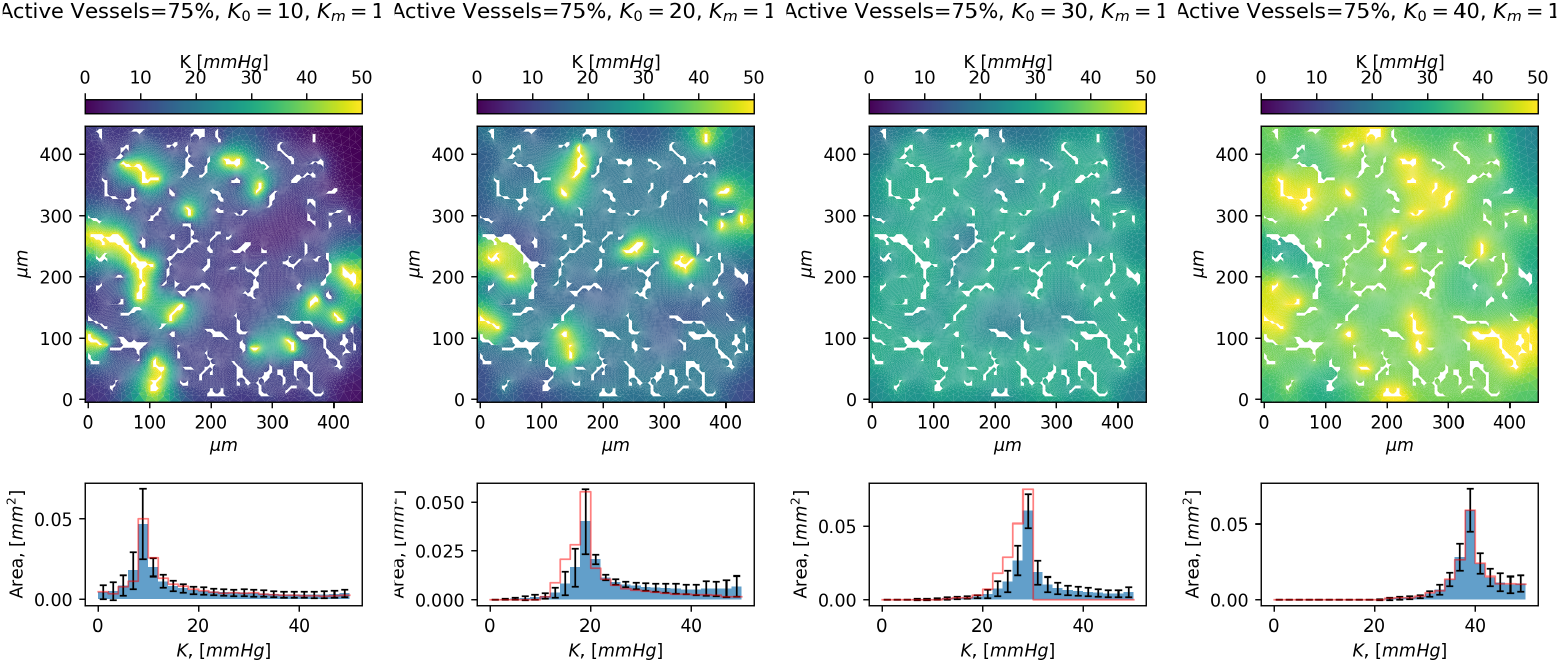
Top: Example oxygen concentration heat maps. Active vessels 75% and the surface *K*_0_ are from left to right: [10, 20, 30, 40]mmHg. Bottom: The mean oxygen concentration histogram is displayed along with standard deviation over the 10 runs.

### Response to irradiation

Based on the oxygen distribution at stationarity, say *K*_∞_(**x**), we can estimate the required irradiation dose *D*_*r*_ needed to ensure a given cell death percentage in the tissue. We evaluate the dose (*D*99) required for cell death of 99%. The cell survival fraction (*S*) around a point **x** depends on the local oxygenation *K*_∞_(**x**). After a dose *D*_*r*_ (compare [9] among others), we assume that it holds

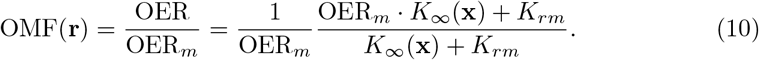

The *α* and *β* values are taken 0.03/*Gy*, and 0.003/*Gy*^2^ respectively and OER_*m*_ = 3, in line with, [7, 9, 11]. To get the dose required for 99% cell death (*D*99) within tumour, we solve numerically *D*_*r*_ from the integral equation

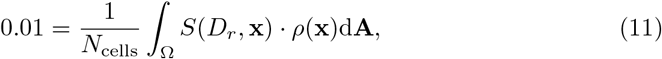

where *ρ*(**x**) is the cell number density (= constant), and *N*_cells_ = ∫_Ω_ *ρ*d**A**. The evaluation of D99 is performed for every of the distributions; the averages and deviations from these averages are computed as well as indicated in Fig. 5.

**Fig 5.**
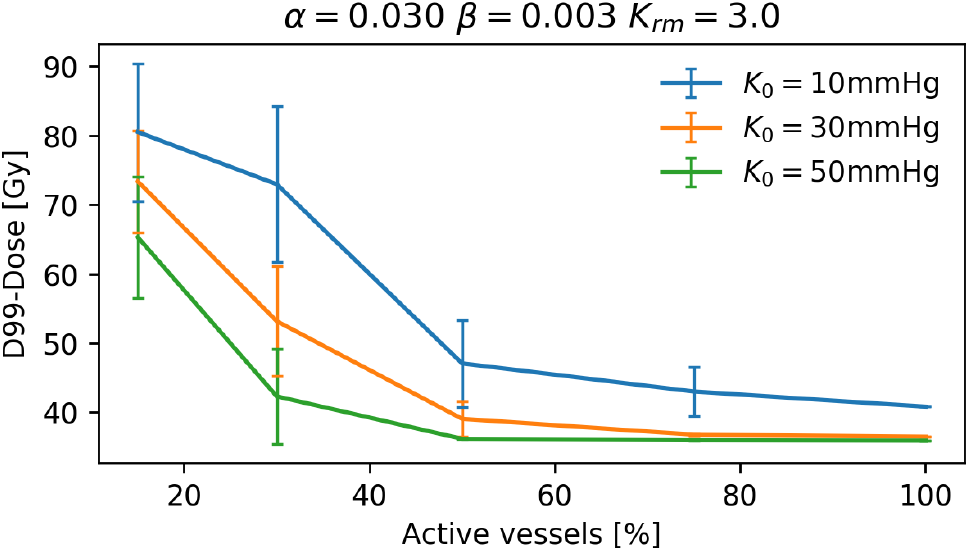
**D99 dose** as a function of active vessel percentage and vessel surface oxygen concentration *K*_0_.

## Discussion. Outlook

One advantage of our approach is that we do not make use of the Green function representation of the stationary distribution of oxygen. Due to the complexity of the tissue structure and due to presence of the vasculature, the exact formula for the corresponding Green function is not available. Furthermore, deviations from “gaussianity” are to be expected. In other words, the precise structure of the Green function for our system would need to be approximated numerically. Instead of doing so, we preferred to deal with the evolution differential equation describing the transport and production of oxygen and capture numerically its effect on the tumour dynamics via irradiation. The ultimate aim is to use mathematical modeling and simulations as well as *in vitro* information to deliver good quantitative estimates of the radiation treatment on the tumour dynamics. To do so, the model needs to be extended with further components (see e.g. [4]). Furthermore, to bring the numerically estimated information at the level of the patient (e.g. to suggest a personalised radiation treatment (dosage, fractionation pattern, time schedule, etc.), we strongly believe that a data-driven modeling needs to improve our predictions. We expect that perhaps MRI scans of the patient’s tumour can bring in such additional information.

There are several ways in which our results could be improved. We mention here only three of the possible directions. However, these are as of now considered out of the scope of this work. Some of these outlets will be considered in forthcoming publications.

- The surface concentrations could be sampled from a given distribution.
- We could use several slices of the tumour for averaging over the resulting oxygen concentration histograms.
- The FEniCS code, approximating with FEM our model (1), can be extended in a straightforward way to handle a 3D computational domain. However, we expect the adaptation of the mesh (for instance via GMSH) to the 3D structure of the blood vessels to be cumbersome.
- The visualisation of the hypoxic regions in 3D requires a special care. The hope is to be able to detect visually the possibility of clogging in the oxygen transport through Ω \ *V*.

## Acknowledgments

We thank O. Richardson (Karlstad, Sweden) as well as G. Powathil (Swansea, UK) for fruitful discussions on the topic.

## Author Contributions

Posing the scientific question and collection of the experimental data: JHL. Modeling of hypoxia: AM. Performed numerical experiments and analysed the data: TK. Wrote the paper: TK, JHL, and AM.

The authors have no conflicts to disclose.

## Footnotes

1 This is only a typical value. We vary it in different ways as reported in the Results section.

2 ”The vessel voxels represents the oxygen level on the surface of the vessel that is available for diffusion into the tissue. In our simulations, we use a default value of 40 mmHg (cf. [7]). However, it is important to note that the model allows for time and space variations in the choice of *K*_0_ - for instance, a direct proportionality with the diameter of the vessel can be incorporated.

3 We point out a feature in the simulation which, with given probability, assigns an active vessel to have *K*_0_ = 50*mmHg* instead of a lower *K*_0_ which is the reference value set to all other active vessels. The probability for such an modification to occur is 1/3 as of now. We proceed this way mainly because the tumours use angiogenesis, which causes vessels to develop in a very unorganised manner, causing large differences in vessel length, trajectory and in tissue vascularisation, hence differences in the oxygen level would be expected.

